# Integrated requirement of non-specific and sequence-specific DNA binding in MYC-driven transcription

**DOI:** 10.1101/2020.05.04.076190

**Authors:** Paola Pellanda, Mattia Dalsass, Marco Filipuzzi, Alessia Loffreda, Alessandro Verrecchia, Virginia Castillo Cano, Mirko Doni, Marco J. Morelli, Marie-Eve Beaulieu, Laura Soucek, Davide Mazza, Marina Mapelli, Theresia Kress, Bruno Amati, Arianna Sabò

**Affiliations:** European Institute of Oncology (IEO) - IRCCS, Via Adamello 16, 20139 Milan, Italy; Center for Genomic Science of IIT@SEMM, Fondazione Istituto Italiano di Tecnologia (IIT), Via Adamello 16, Milan 20139, Italy; Experimental Imaging Center, IRCCS San Raffaele Scientific Institute, Milan, Italy; Peptomyc S.L., Edifici Cellex, Barcelona 08035, Spain; Vall d’Hebron Institute of Oncology (VHIO), Edifici Cellex, Barcelona 08035, Spain; Institució Catalana de Recerca i Estudis Avançats (ICREA), Barcelona 08010, Spain; Department of Biochemistry and Molecular Biology, Universitat Autònoma de Barcelona, Bellaterra 08193, Spain

**Keywords:** Myc, DNA binding, E-box, promoter, transcription

## Abstract

Eukaryotic transcription factors recognize specific DNA sequence motifs, but are also endowed with generic, non-specific DNA-binding activity: how these binding modes are integrated to determine select transcriptional outputs remains unresolved. We designed mutants of the MYC transcription factor bearing substitutions in residues that contact either the DNA backbone or specific bases within the consensus binding motif (E-box), and profiled their DNA-binding and gene-regulatory activities in murine cells. Our data reveal that non-specific DNA binding is required for MYC to engage onto active regulatory elements in the genome, preceding sequence recognition; beyond merely stabilizing MYC onto select target loci, sequence-specific binding contributes to its precise positioning and – most unexpectedly – to transcriptional activation *per se*. In particular, at any given binding intensity, promoters targeted via the cognate DNA motif were more frequently activated by MYC. Hence, seemingly promiscuous chromatin interaction profiles actually encompass diverse DNA-binding modalities, driving defined, sequence-dependent transcriptional responses.

## Introduction

The transcription factor Myc orchestrates complex gene expression programs that foster cell growth and proliferation, in both normal and cancer cells (Kress et al., 2016, Kress et al., 2015, Muhar et al., 2018, Sabò et al., 2014, Tesi et al., 2019, Walz et al., 2014). Myc dimerizes with Max (Blackwood & Eisenman, 1991) to bind DNA with a preference for the E-box consensus sequence CACGTG (Blackwell et al., 1993, Solomon et al., 1993), through which it activates transcription (Amati et al., 1992, Kretzner et al., 1992). Within cells, however, Myc promiscuously associates with active chromatin (Guccione et al., 2006, Kim et al., 2008, Soufi et al., 2012), owing most likely to a combination of general accessibility (Sabò et al., 2014), protein-protein interactions (Richart et al., 2016, Thomas et al., 2019, Thomas et al., 2015) and non-specific DNA binding (Brownlie et al., 1997, Ferre-D’Amare et al., 1993, Nair & Burley, 2003, Sauvé et al., 2007): consequently, when expressed at high levels, Myc can be detected on virtually all active promoters and enhancers in the genome (Guo et al., 2014, Lin et al., 2012, Nie et al., 2012, Sabò & Amati, 2014, Sabò et al., 2014, Walz et al., 2014). Hence, while Myc-activated promoters tend to show overrepresentation of E-box motifs (Tesi et al., 2019) and stronger Myc recruitment (de Pretis et al., 2017, Lorenzin et al., 2016, Walz et al., 2014), the role of DNA sequence recognition in Myc activity remains to be clarified.

Here, we exploited the structure of the DNA-bound Myc/Max dimer (Nair & Burley, 2003) to design Myc mutants lacking either non-specific contacts with the DNA phosphodiester backbone (R364/366/367-A; Myc^RA^) or base-specific interactions within the E-box consensus (H359/E363-A; Myc^HEA^). When ectopically expressed in cells, Myc^RA^ showed pervasive loss of genome interactions and transcriptional activity, associated with increased intra-nuclear mobility. Myc^HEA^ instead retained DNA binding and mobility profiles comparable to those of the wild-type protein, but failed to recognize the E-box, and could not activate Myc-target genes. Concurrently, Myc^HEA^ gained weak affinity for an alternative motif, driving aberrant activation of genes not normally regulated by Myc. Altogether, non-specific DNA binding is essential for Myc to engage onto genomic regulatory regions, but does not *per se* support transcriptional activation; sequence recognition, instead, contributes to activation at three distinct levels: stabilization and positioning of the factor onto specific DNA motifs, and promotion of its transcriptional activity.

## Results

### Structure-based mutagenesis of the Myc DNA-binding domain

Myc/Max dimerization depends upon the contiguous helix-loop-helix and leucine zipper domains of each protein (HLH-LZ: ca. 70 amino acids) and is a strict pre-requisite for DNA binding, mediated by the short basic region (15 a.a.) that precedes the HLH (Amati et al., 1992, Blackwood & Eisenman, 1991). In line with those biochemical findings, structural studies on DNA-bound or free dimers, including Myc/Max (Nair & Burley, 2003, Sammak et al., 2019), Max/Max (Brownlie et al., 1997, Ferre-D’Amare et al., 1993, Sauvé et al., 2007, Sauvé et al., 2004) and other bHLH proteins (Murre, 2019), showed that dimerization allows positioning of the basic regions for insertion into the DNA major groove (**Fig. EV1A**).

We targeted two groups of residues involved in DNA contacts within the Myc basic region (**Fig. EV1B**). First, R364, R366 and R367 interact with the phosphodiester backbone. Early data showed that a mutant with the triple alanine substitution (hereafter Myc^RA^) was proficient in dimerization with Max, but could not activate an E-box-driven reporter gene (Amati et al., 1992): here, we further characterized this mutant as a candidate for loss of generic (non-sequence specific) DNA binding. Second, H359 and E363 form H-bonds with the invariant bases of the E-box consensus (<u>CA</u>NN<u>TG</u>): we thus substituted these residues with alanines to impair sequence-specific recognition, (Myc^HEA^).

When introduced by transfection in HEK-293T cells, Myc^RA^ and Myc^HEA^ were expressed and co-precipitated endogenous Max as efficiently as Myc^WT^ (**Fig. EV1C**). Polypeptides spanning the Myc and Max bHLH-LZ domains were purified from *E. coli* and used to test binding to a fluorescently labelled E-box DNA probe *in vitro*: as expected (Beaulieu et al., 2012), combining Max with Myc^WT^ yielded a distinct protein-DNA complex compared to the Max homodimer, while this occurred neither with Myc^HEA^ nor Myc^RA^ (**Fig. EV1D**). Hence, at this level of resolution, both of the Myc mutants associated with Max but the resulting dimers were unable to bind the E-box.

### Differential impairment in genome recognition by Myc^HEA^ and Myc^RA^

To address the ability of our Myc mutants to interact with genomic DNA *in vivo*, we expressed them in 3T9 fibroblasts as 4-hydroxytamoxifen (OHT)-dependent MycER^T2^ chimeras (MycER^WT^, MycER^RA^ and MycER^HEA^: **Fig. EV2A**). We thus treated the cells with OHT for 4h and profiled them by ChIP-seq with antibodies recognizing either Myc or the ER moiety: while the former could not discriminate between the endogenous and exogenous forms, the latter detected only the MycER variants, allowing us to specifically follow the mutated proteins (**Fig. EV2B**). As previously reported (de Pretis et al., 2017, Sabò et al., 2014), MycER^WT^ showed widespread association with active regulatory elements (i.e. promoters and enhancers) throughout the genome of 3T9 cells, as defined by the presence of active histone marks (H3K4me1, H3K4me3, H3K27ac), RNA Polymerase II (RNAPII) and DNAseI hypersensitivity (**Fig. EV2C**): this effect - sometimes termed “invasion” (Lin et al., 2012, Sabò et al., 2014) - was greatly attenuated with MycER^RA^ and instead strengthened with MycER^HEA^, which showed even wider spreading than MycER^WT^ onto active chromatin, in particular at gene-distal regions. Consistent with these profiles, peak-calling identified 16,762 peaks for MycER^WT^, 5,615 (33%) for MycER^RA^ and 23,873 (142%) for MycER^HEA^.

To better characterize the effects of the mutations on DNA binding, we focused our attention on the occurrence of consensus elements at MycER-binding sites, considering the canonical E-box CACGTG (or #1) and four previously identified variants (#2-5: CACG<u>C</u>G, CA<u>T</u>G<u>C</u>G, CACG<u>A</u>G, CAC<u>A</u>TG) (Allevato et al., 2017, Blackwell et al., 1993, Grandori et al., 1996, Guo et al., 2014, Perna et al., 2012). As previously observed (Guccione et al., 2006, Kim et al., 2008, Sabò & Amati, 2014, Soufi et al., 2012), binding hierarchy correlated primarily with chromatin, rather than with the presence of these consensus motifs (**Fig. 1A**). This notwithstanding, the motifs significantly contributed to the MycER^WT^ profiles, as evidenced by three distinctive features: first, the percentage of MycER^WT^ peaks containing at least one motif within ±100bp from the peak summit was significantly above background frequency (empty vector: EV; **Fig. 1B**); second, MycER^WT^ peaks with canonical E-boxes showed stronger average intensities, followed by those with variant motifs, and ultimately by motif-free peaks (**Fig. 1C**); third, the DNA motifs were most frequently centered under the peak summit, implying that they contributed to the precise positioning of MycER^WT^ (**Fig. 1D**). Most importantly, both of the MycER mutants showed substantial loss of those sequence-associated features (**Fig. 1B-D**) – note that while MycER^HEA^ retained slightly higher average intensities in the presence of E-boxes (**Fig. 1C**), this might be due to residual recognition of half-sites by MycHEA/Max dimers (see below).

**Figure 1.**
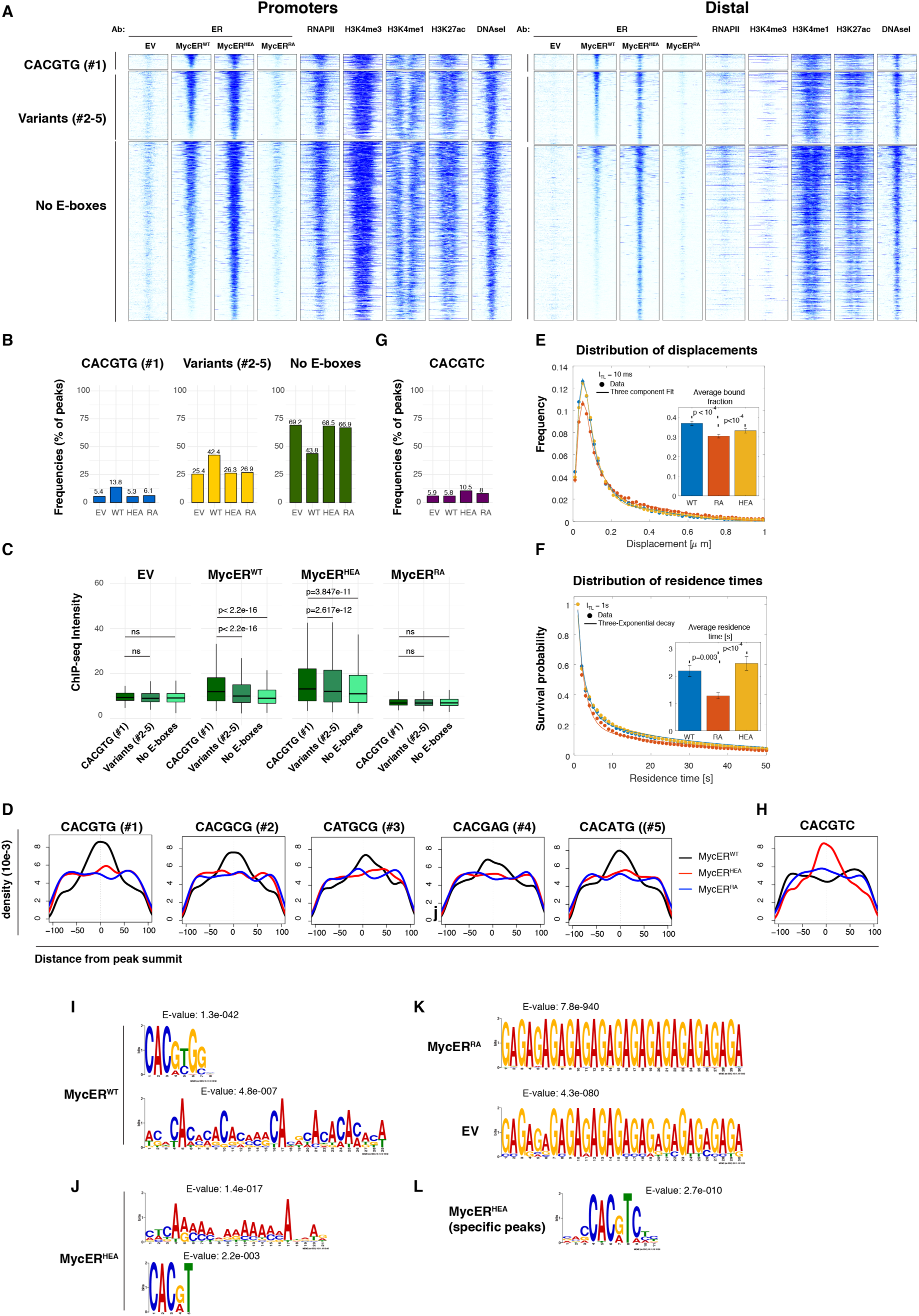
Myc^HEA^ and Myc^RA^ show differentially altered genome-binding profiles. 3T9 fibroblasts transduced with MycER^WT^-, MycER^HEA^-, MycER^RA^-expressing retroviruses or the control empty vector (EV) were treated with OHT (4h) and profiled by ChIP-seq with anti-ER antibodies. **A** Heatmaps representing normalized ChIP-seq intensities at MycER-associated promoters or distal sites, as indicated. Each row represents a genomic site called in at least one of the experimental samples, with each column spanning a 4 kb-wide genomic interval centered on the union of MycER peaks. All sites are ranked according to the intensity of the MycER^WT^ signal, and divided based on the presence of the indicated DNA motifs (#1, #2-5 or none) in an interval of ±100 bp around their peak summit. The data for RNAPII, histone marks (H3K4me3, H3K4me1, H3K27ac) and DNAseI hypersensitivity are from 3T9-MycER^WT^ fibroblasts without OHT (Sabò et al., 2014). **B** Frequency of peaks (as %) that contain the indicated motif within ±100 bp from the peak summit in each ChIP-seq sample (EV, WT, HEA and RA). **C** Average ChIP-seq intensities for regions containing the indicated motifs in each sample. p-values were calculated using Wilcoxon’s test. **D** Density plots showing the distribution of the indicated motifs in a ±100 bp interval from the peak summit. **E** Single molecule tracking at high frame rate: the time of *t*_*tl*_ = 10ms between two images allows to estimate the distribution of displacements, that is then fit by a three-component diffusion model to estimate the fraction of molecules immobilized on chromatin (Average bound fraction, inset, see Fig. EV4 and Supplementary Methods). Note that Myc^RA^ displays a significantly lower bound fraction than Myc^WT^ and Myc^HEA^ (*n*_*cells*_= 34, 30 and 35, and *n*_*displacements*_ = 78550, 59783 and 83801 for Myc^WT^, Myc^RA^ and Myc^HEA^, respectively). Error-bar: SD. Statistical significance evaluated by permutation tests. **F** Single molecule tracking at lower frame rate (*t*_*tl*_spanning between 200 ms and 2s) allows to quantify the distribution of residence times (i.e. the duration of binding events), revealing a significantly shorter average for Myc^RA^, relative to either Myc^WT^ or Myc^HEA^ (*n*_*cells*_= 35, 35 and 31, and *n*_*Bound–molecules*_ = 2452, 2084 and 2171 for Myc^WT^, Myc^RA^ and Myc^HEA^, respectively). Error-Bar: SD. Statistical test: Anova-Tukey. **G, H** As in (B) and (D), respectively, for the CACGTC motif. **I**-**K** *De novo* motif discovery analysis performed underneath the summit of the top 200 peaks called with (**I**) MycER^WT^, **(J**) MycER^HEA^, (**K**) MycER^RA^ and empty vector (EV) infected cells. The position weight matrixes of predicted DNA binding motifs are shown together with their E-values. **L** As in (I) for the top 200 MycER^HEA^ -specific peaks (i.e. not bound by MycER^WT^).

The above results were obtained with over-expressed proteins. In this regard, it must be noted that ChIP-seq profiles represent the sum of sequence-specific and non-specific binding events in large cell populations: we thus predicted that at reduced protein levels – i.e. when weak binding events should be the least detectable over background – Myc^HEA^ should show deeper binding defects relative to Myc^WT^. To address this prediction, we used mouse fibroblasts expressing a doxycycline-dependent tet-Myc transgene (cb9 cells) and engineered the endogenous c-*myc* locus in two different manners: first, we introduced the HEA mutation into both c-*myc* alleles (cb9-*myc*^HEA^; see methods); second, we inactivated the endogenous product (cb9^Δmyc^; **Fig. EV3A**) and introduced either Myc^WT^, Myc^HEA^ or Myc^RA^ by retroviral transduction. Shut down of tet-Myc (-dox) thus left only the full-length Myc^WT^, Myc^HEA^ or Myc^RA^ proteins, expressed from either the exogenous or endogenous genes (**Fig. EV3B**). By ChIP-seq profiling, the over-expressed Myc^HEA^ protein (but not Myc^RA^) showed widespread DNA binding, while cb9-*myc*^HEA^ cells showed substantial loss binding relative to the control cb9-*myc*^WT^ clones, confirming that protein levels directly impacted the detection of Myc^HEA^ on chromatin (**Fig. EV3C**). Most importantly Myc^HEA^ showed a loss of E-box selectivity analogous to that described for MycER^HEA^ (**Fig. EV3D-H**).

Altogether, Myc^HEA^ lacks E-box recognition but retains the propensity to distribute along active chromatin when overexpressed, consistent with the notion that chromatin features (i.e. accessibility, composition, protein-protein interactions, etc…) rather than DNA sequence are the primary determinants of Myc binding (Guccione et al., 2006, Kim et al., 2008, Richart et al., 2016, Sabò & Amati, 2014, Soufi et al., 2012, Thomas et al., 2019, Thomas et al., 2015). However, binding must also rely upon close contacts with the DNA backbone, as demonstrated by the loss of interaction seen with the Myc^RA^ mutant. We surmise that non-specific DNA binding may contribute to tether Myc onto active regulatory regions (promoters and enhancers), restricting its free diffusion in the nucleoplasm and allowing local scanning of the DNA sequence.

As a corollary, loss of this initial tethering step in Myc^RA^ - but not Myc^HEA^ - would be expected to cause an increase in protein mobility relative to Myc^WT^. To address this issue, we used single-molecule tracking microscopy (Gebhardt et al., 2013, Mazza et al., 2012) on live mouse fibroblasts expressing Myc-HaloTag fusion proteins: indeed, Myc^HEA^ behaved as Myc^WT^, while Myc^RA^ underwent significant gains in mobility, showing both a reduced proportion of immobilized molecules (**Fig. EV4A,B; Fig. 1E**) and shorter average residence times on chromatin (**Fig. EV4C,D; Fig. 1F**).

### Myc^HEA^ gains recognition of an alternative non-E-box motif

To better characterize the alterations in DNA sequence recognition caused by the HEA and RA mutations, we performed *de novo* motif analysis on the top 200 sites bound by each MycER variant, considering either all ChIP-seq peaks (**Fig. 1I-L**) or promoter-associated and distal peaks separately, to account for differences in base composition (**Appendix Figure S1**). As expected, MycER^WT^ peaks enriched with high statistical significance for position weight matrices (PWMs) matching either the canonical E-box CACGTG or variants #2-5 (**Fig. 1I, Appendix Figure S1A**), but also for degenerate AC-rich motifs, which incidentally included several CAC half-sites. Remarkably, MycER^HEA^ lost the main PWM of MycER^WT^ but still enriched for the AC-rich motifs, and secondly – with lower significance – for the partial E-box motifs CAC(^G^/_A_)TN or CACG(^C^/_T_)C (**Fig. 1J, Appendix Figure S1B**), consistent with the CAC half-site being contacted by the wild-type Max moiety (**Fig. EV1A,B**). MycER^RA^ peaks instead were not enriched for specific motifs over the non-specific background detected in control cells (EV; **Fig. 1K**).

As noted above, MycER^WT^ and MycER^HEA^ showed largely superimposable binding profiles, owing most likely to general accessibility: restricting our motif analysis to sites bound only by MycER^HEA^ led to improved definition of the aforementioned partial motifs to CAC(^G^/_A_)TC (**Fig. 1L, Appendix Figure S1C**). Thus, consistent with the contacts established by residues H359 and E363 with the conserved G6-C1’ base-pair (CACGTG: **Fig. EV1B**), substitution of these amino-acids not only impaired E-box recognition, but also altered the specificity to CACGTC: this new motif was specifically enriched (10-15% of the peaks) and precisely positioned with the HEA, but not the WT or RA proteins (**Fig. 1G,H; Fig. EV3G,H**). It is important to note, however, that this did not amount to a full subversion of DNA-binding specificity, since (*i.*) MycER^HEA^-only sites - which allowed the most stringent definition of the alternative motif - were bound at weaker levels than those shared with MycER^WT^ (**Fig. EV2C**); (*ii.*) when considering all sites, CACGTC was enriched by MycER^HEA^ only after the degenerate AC-rich motifs, while MycER^WT^ enriched primarily for the canonical E-box (**Fig. 1I,J; Appendix Figure S1A,B**); (*iii.*) Myc^HEA^ knock-in cells showed extensive loss of DNA-binding activity (**Fig. EV3C**).

### Sequence recognition determines transcriptional activation

To address the impact of DNA-binding alterations on transcriptional activity, we established RNA-seq profiles following OHT treatment of 3T9-MycER^WT^, MycER^HEA^ and MycER^RA^ cells. As previously observed (de Pretis et al., 2017, Sabò et al., 2014), MycER^WT^ elicited the up- and down-regulation of equivalent numbers of genes (ca. 1000 at 4h, 2000 at 8h): while this effect was largely lost with the MycER^RA^ mutant, MycER^HEA^ mobilized even more mRNAs (**Fig. 2A**) but with profiles entirely different from those of MycER^WT^ (**Fig. 2B,C)**. In particular, focusing on MycER^WT^-regulated genes revealed that MycER^HEA^ modulated their expression inconsistently, with a continuum of effects ranging from activation to repression (**Fig. 2D)**. Moreover, MycER^HEA^ regulated additional genes, not modulated by MycER^WT^ (**Fig. 2E)**. Closer scrutiny revealed that the differences between MycER^WT^- and MycER^HEA^ -regulated transcriptional programs were attributable to DNA sequence: in particular, genes activated by either one or the other form of MycER (or both) showed increased frequency of the cognate consensus motifs under the corresponding ChIP-seq peak in the promoter (**Fig. 2F**).

**Figure 2.**
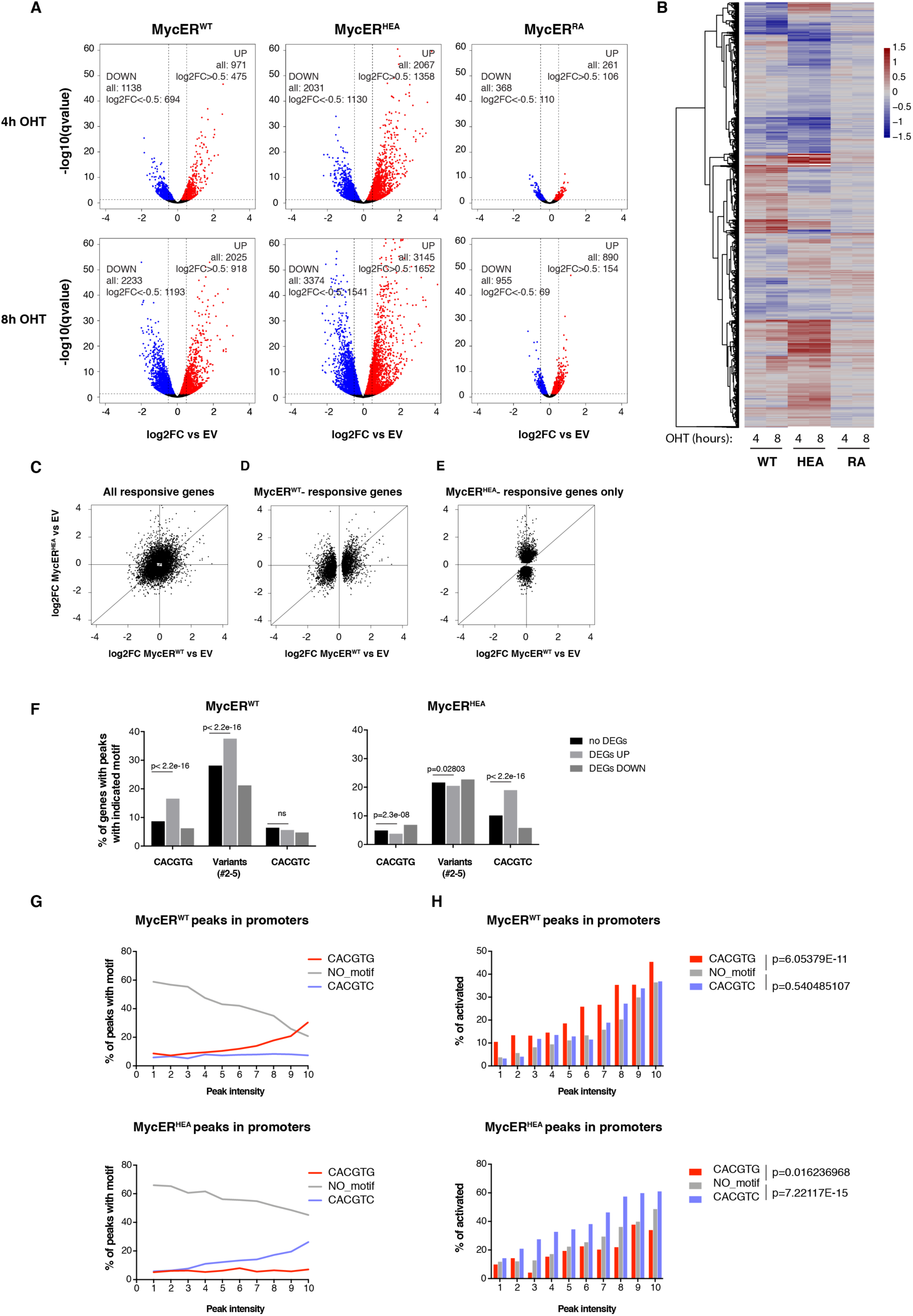
DNA sequence recognition determines transcriptional regulation. 3T9 fibroblasts expressing the indicated MycER proteins and control cells (EV) were treated with OHT (4h, 8h) and profiled by RNA-seq. **A** Fold-change of each annotated mRNA (log2FC, relative to the EV control), plotted against its q-value (-log10). mRNAs showing significant up- and down-regulation (qval<0.05) are marked in red and blue, respectively, and their numbers reported in the graphs, along with those with qval<0.05 and|log2FC|>0.5. **B** Heatmap representing the same log2FC values as in a. (restricted to those mRNAs with qval<0.05 in at least one of the MycER samples). **C**-**E** Scatter plots confronting fold-change values (defined as in A) in response to MycER^WT^ (x-axis) and MycER^HEA^ (y-axis), including either all of the mRNAs called as DEGs (qval<0.05) in at least one of the samples (C), all MycER^WT^-regulated DEGs (D), or the MycER^HEA^-specific DEGs (E). **F** Percentage of promoters with the indicated DNA motifs under the ChIP-seq peak (±100 bp from the peak summit) within each regulatory class (no DEG, UP or DOWN) for either MycER^WT^ (left) or MycER^HEA^ (right). p-values calculated with Fisher’s exact test. **G** Percentage of promoter-associated ChIP-seq peaks with the indicated motifs, as a function of peak intensity (binned in deciles: 1-10) for either MycER^WT^ (top) or MycER^HEA^ (bottom). **H** Percentage of DEG UP genes (qval<0.05) as a function of peak intensity (as in G). Statistical test: Chi-squared against the “no motif” condition, performed on the entire series

Two non-mutually exclusive mechanisms may underlie the connection between sequence recognition and transcriptional activation. First, the presence of the cognate binding motif stabilizes DNA binding by either MycER^WT^ or MycER^HEA^, as evidenced by peak intensities in ChIP-seq profiles (**Fig. 2G**): as a consequence, the extended residence time of the transcription factor on DNA may increase the probability of activation. In line with this scenario, Myc-induced transcriptional programs in diverse cell types correlated with the relative gain in Myc binding at promoters (de Pretis et al., 2017, Lorenzin et al., 2016, Tesi et al., 2019, Walz et al., 2014), as confirmed in our experiments (**Fig. S6A**): moreover, and consistent with the proposed mechanism, DNA binding and activation by either form of MycER were associated with enrichment of the cognate DNA motif (**Fig. S6B**). Second, beyond residence time, sequence recognition may directly contribute to the molecular activity of the transcription factor. Remarkably, our data also provided support for this scenario: indeed, at any given binding intensity (bins 1-10), loci targeted via the cognate DNA motif were more frequently activated by either MycER^WT^ or MycER^HEA^, the opposite motif serving as negative control (**Fig. 2H**).

Unlike activated genes, those down-regulated by either MycER^WT^ or MycER^HEA^ recruited the transcription factor with the lowest efficiency and lacked enrichment of the cognate binding motif **(Fig. EV5A,B)**: hence, as previously proposed (Baluapuri et al., 2019, de Pretis et al., 2017), repression by either MycER^WT^ or MycER^HEA^ may be largely indirect. Moreover, MycER^HEA^-repressed loci included known Myc-dependent genes (Lorenzin et al., 2016, Muhar et al., 2018, Perna et al., 2012, Tesi et al., 2019) (**Fig. EV5C**): we surmise that by binding these loci without being able to recognize the E-box, MycER^HEA^ may impair their sustained activation by endogenous Myc.

Altogether, our data show that sequence recognition is essential to establish adequate Myc-dependent transcriptional programs: most importantly, this step is subsequent to engagement of the factor on active chromatin, mediated by non-specific DNA binding.

### Myc^RA^ and Myc^HEA^ are unable to sustain cell proliferation

In order to assess the biological activity of our Myc mutants, we monitored their ability to support cell proliferation upon tet-Myc shutdown in cb9^Δmyc^ cells: while Myc^WT^, Myc^HEA^ and Myc^RA^ were expressed at similar levels in those conditions (**Fig. EV3B**), only Myc^WT^ supported colony formation, cell proliferation and DNA-synthesis (**Fig. 3A-C**; -dox). When the tet-Myc transgene was maintained active, Myc^HEA^-expressing cells showed reduced proliferative activity (**Fig. 3A-C**; +dox), with similar effects upon activation of MycER^HEA^ in 3T9 cells (**Fig. EV2D**). This growth-inhibitory activity depended upon over-expression of Myc^HEA^, as the knock-in cb9-*myc*^HEA^ clones showed no proliferative defects relative to their WT counterparts in the presence of tet-Myc (**Fig. 3D**; +dox). Finally when expressed in the c-*myc*-null rat fibroblast cell line HO15.19 (Mateyak et al., 1997) (**Fig. 3E**), Myc^WT^ promoted proliferation and colony formation to levels close to those of parental TGR1 cells, while Myc^HEA^ and Myc^RA^ showed no effect (**Fig. 3F, G**). In summary, neither Myc^HEA^ nor Myc^RA^ could compensate for the loss of endogenous Myc, and Myc^HEA^ had dominant-negative activity, manifest only upon over-expression in Myc-proficient cells. Most noteworthy here, mutations in the residue equivalent to Myc-E363 in other bHLH proteins also conferred DN activity, associated with altered DNA binding (Boisson et al., 2013, Luchtel et al., 2019, Marchegiani et al., 2015).

**Figure 3.**
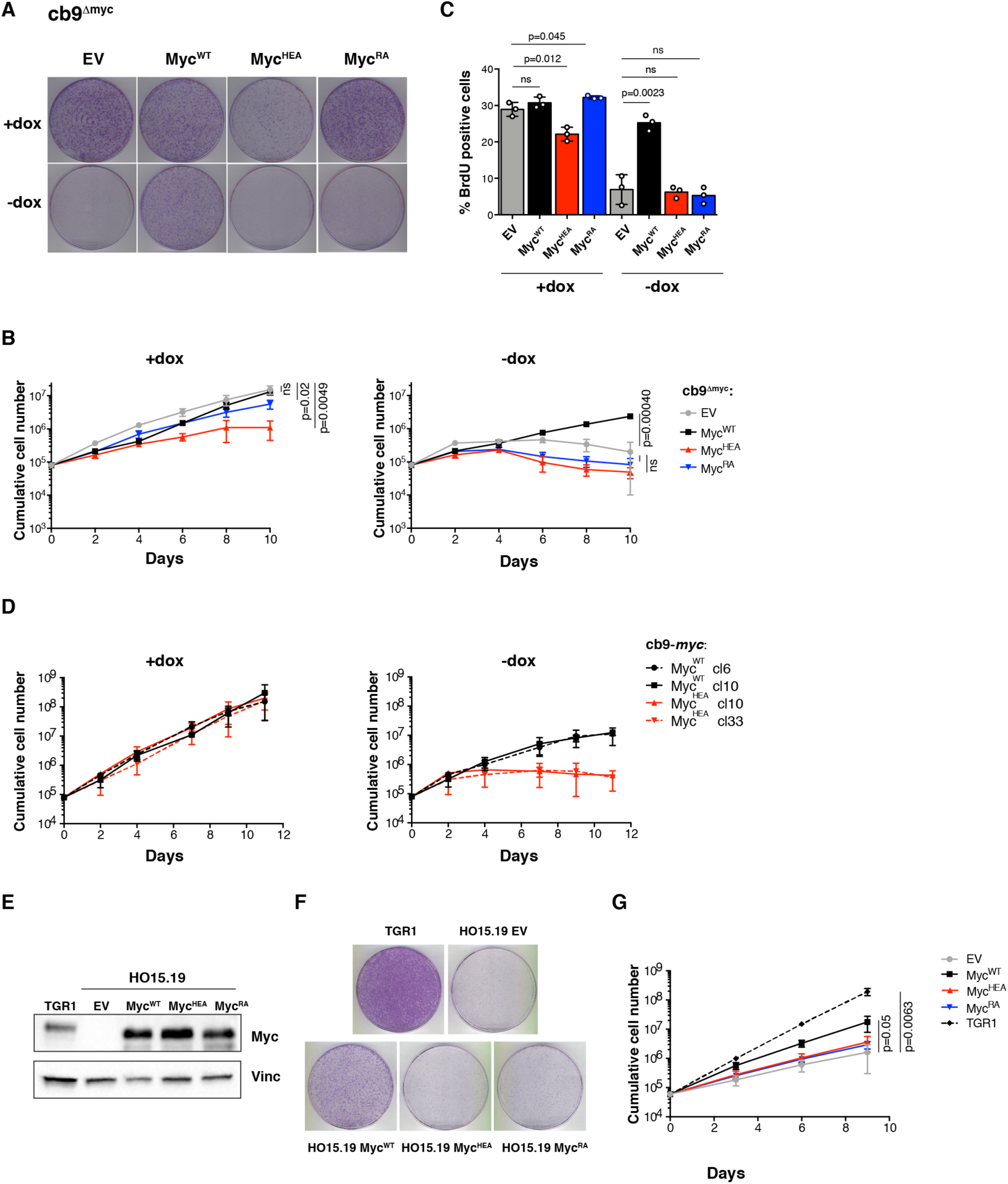
Myc^HEA^ and Myc^RA^ are defective in sustaining cell proliferation. **A** Colony formation for cb9^Δmyc^ cells infected with retroviral vectors expressing the indicated Myc proteins; all cells were expanded with doxycycline prior to the final plating step, upon which the compound was either maintained (+dox), or removed (-dox) to switch off the tet-Myc transgene. **B** Cumulative cell counts for cb9^Δmyc^ cells upon serial passaging with or without dox (removed at time day 2). **C** Percentage of BrdU positive cb9^Δmyc^ cells, 24h after dox removal. **D** As in (B) for cb9-*myc*^WT^ and *myc*^HEA^ cells clones. **E** Immunoblot analysis of c*-myc*^*-/-*^ HO15.19 rat fibroblasts infected with retroviral vectors expressing the indicated Myc proteins. Parental TGR1 cells serve as wild-type control. **F** Colony formation for the same cells as in (E). **G** cumulative cell counts for the same cells as in (E).

## Discussion

Unlike pioneer factors that can access DNA in closed chromatin (Kim et al., 2008, Soufi et al., 2012), Myc and other bHLH proteins depend upon a pre-existing active chromatin state (Guccione et al., 2006, Guo et al., 2014, Kim et al., 2008, Sabò et al., 2014, Soufi et al., 2012, Xin & Rohs, 2018). Moreover, while recognizing specific DNA sequence motifs, all of these transcription factors are also endowed with generic, non-specific DNA-binding activities: how these features are integrated to determine genomic binding profiles and transcriptional outputs remains largely unresolved. We have addressed this question for Myc by mutating residues that contact either the DNA backbone (Myc^RA^) or specific bases within the E-box consensus motif (Myc^HEA^). Expression of these mutants in cultured mouse fibroblasts allowed us to dissect their DNA-binding and gene-regulatory properties, unravelling several key features. First, besides open chromatin, non-specific DNA binding is required for Myc to engage on genomic regulatory regions, as a prerequisite for sequence-specific recognition; most importantly, this initial step underlies the majority of the cross-linking events detected by ChIP-seq in cells with high Myc levels, emphasizing the need to discriminate between specific and non-specific DNA binding. Second, DNA sequence recognition contributes not only to stabilization of the Myc protein onto select DNA motifs, but also to its transcriptional activity *per se*, implying some form of communication between the C-terminal bHLH-LZ and the N-terminal transactivation domain of Myc (Amati et al., 1992, Barrett et al., 1992). For example, sequence-specific binding may elicit allosteric changes that modulate transcriptional activity, as proposed for or other TFs such as the glucocorticoid receptor (Watson et al., 2013) or the bHLH protein MyoD (Huang et al., 1998); in line with this concept, Max bHLH-LZ homodimers showed subtle structural differences when bound to specific vs. non-specific DNA (Sauvé et al., 2007), but whether the same occurs with Myc/Max remains unknown. Notwithstanding its molecular underpinning, this unexpected connection reveals that widespread ChIP-seq profiles actually include different binding modalities, which in turn drive specific transcriptional responses (Kress et al., 2015, Sabò & Amati, 2014, Sabò et al., 2014) rather than a general increase in transcriptional activity (Lin et al., 2012, Nie et al., 2012). Finally, besides transcriptional regulation *per se*, Myc was suggested to act in the resolution of transcription- and replication-associated stresses (Baluapuri et al., 2020): whether these additional effects of Myc involve DNA sequence recognition remains to be addressed.

## Materials and Methods

### MYC mutagenesis and subcloning

Mutagenesis of His 359 and Glu 363 to Alanine in human *MYC* (Myc^HEA^ mutant) was performed using the QuikChange Site-Directed Mutagenesis Kit (Agilent Technologies # 200519), according to the manufacturer’s instructions, with a pBabe-hygro plasmid (BH) (Morgenstern & Land, 1990) containing the human *MYC* cDNA (Amati et al., 1992) as template. The resulting pBH-Myc^HEA^ plasmid was then sequenced to verify the correctness of the whole mutated cDNA (codon 359: CAC to GCC; codon 363: GAG to GCG). The resulting mutant cDNA and that encoding the Arg 364/366/367 to Ala mutant (Amati et al., 1992) (here Myc^RA^) were subcloned in the retroviral vector pQCXIH (Clontech) or in the transfection vector pCMV-FLAG. For expression as 4-hydroxy-tamoxifen (OHT)-dependent MycER^T2^ chimeras (Littlewood et al., 1995), the full-length *MYC* cDNAs were fused in-frame upstream of the variant estrogen receptor hormone-binding domain (ER^T2^) into the retroviral vector pBabe-puro (BP) (Morgenstern & Land, 1990).

Myc-HaloTag vectors were generated using the Gibson Assembly Protocol (Gibson et al., 2009): HaloTag and *MYC* cDNAs (encoding Myc^WT^, Myc^HEA^ or Myc^RA^) were PCR-amplified with primers containing overlapping sequences, and combined with the BP vector (digested with BamHI and EcoRI) in the Gibson Assembly Reaction, to create the final BP-Myc-HaloTag plasmids with in-frame Myc-HaloTag fusions.

### Cell lines

The cb9 tet-Myc cell line was produced through the 3T3-immortalization protocol starting from mouse embryonic fibroblasts (E14.5) obtained from Rosa26-rtTA/tet-Myc mice (Croci et al., 2017) and used to derive the cb9^Δmyc^ and the cb9-*myc*^HEA^ cell lines (see below). All the cell lines used in this work were grown in DMEM, supplemented with 10% fetal bovine serum, 2 mM L-glutamine and 1% penicillin/streptomycin. For the cb9^Δmyc^ and the cb9-*myc*^HEA^ cell lines, medium was also complemented with doxycycline (1 µg/ml) to keep the tet-Myc transgene expressed, unless otherwise specified. Mouse 3T9 fibroblasts were infected with BP-MycER^WT^, MycER^HEA^ or MycER^RA^ retroviruses, and selected for 2 days with puromycin (1.5 μg/ml); activation of the MycER fusion proteins was achieved addition of OHT to the culture medium (400 nM). Rat HO15.19 cells (Mateyak et al., 1997) and mouse cb9^Δmyc^ cells were infected with BH- and QCXIH-based recombinant retroviruses, respectively, expressing either Myc^WT^, Myc^HEA^ or Myc^RA^; infected cells were selected with hygromycin (150 µg/ml) for 4 days.

### Western Blot and co-immunoprecipitation

For western blot, protein extraction was performed by resuspending the cells in lysis buffer (300 mM NaCl, 1% NP-40, 50 mM Tris-HCl pH 8.0, 1 mM EDTA) freshly supplemented with protease inhibitors (cOmplete™, Mini Protease Inhibitor Cocktail, Roche-Merck, #11836153001), followed by brief sonication. After centrifugation at 16000g for 15 minutes at 4°C, cell extracts were quantified with the Bradford-based Protein Assay kit (Bio-Rad Protein Assay, #5000006). After addition of 6X Laemmli buffer (375 mM Tris-HCl, 9% SDS, 50% glycerol, 9% beta-mercatoethanol and 0.03% bromophenol blue), lysates were boiled for 5 minutes, electrophoresed on SDS-PAGE gels (7.5% Polyacrylamide), transferred onto nitrocellulose membranes and protein expression detected with the indicated primary antibodies (see below). Chemiluminescence was detected using a CCD camera (ChemiDoc XRS+ System, Bio-Rad). Quantification of protein levels was performed using the Image Lab software (Bio-Rad, version 4.0).

For co-immunoprecipitation experiments, 293T cells were transfected overnight with calcium phosphate with 5 µg of plasmids encoding FLAG-tagged Myc^WT^, Myc^HEA^, Myc^RA^ or EV (empty vector) and collected 48h after transfection. After two washes in ice-cold PBS, cells were scraped in 4 ml of ice-cold NHEN buffer (20 mM Hepes pH 7.5, 150 nM NaCl, 0.5% NP-40, 10% glycerol, 1 mM EDTA) freshly supplemented with protease inhibitors (cOmplete™, Mini Protease Inhibitor Cocktail, Roche-Merck, #11836153001) and lysed for 20 minutes on a rotating wheel at 4°C. Complete cell disruption and DNA fragmentation was performed with three cycles of sonication (30 seconds on, 30 seconds off) with a Branson Sonifier 250 (Output Control = 2) equipped with a 3.2 mm Tip (Branson, #101-148-063). Lysates were cleared by centrifugation at 16000g for 15 minutes at 4°C, and protein concentration determined with the Bradford-based Protein Assay kit (Bio-Rad Protein Assay, #5000006). The immunoprecipitation of FLAG-Myc was performed by incubating 2 mg of cell lysate with 40 µl of Anti-FLAG M2 affinity gel (Sigma-Aldrich #A2220) for 3h in a final volume of 1 ml of NHEN buffer with agitation at 4°C. The beads were then washed five times with 1 ml of wash buffer (20 mM Hepes pH 7.5, 150 nM NaCl, 0.1% Tween, 10% glycerol, 1 mM EDTA), resuspended in 1X Laemmli buffer and boiled for 10 minutes. In parallel, 2.5% of the material used for the IP was collected to be loaded as input.

### Production and purification of Myc and Max bHLH-LZ peptides, and DNA-binding assay

Peptides spanning the bHLH-LZ domains of Max (p21 isoform; sequence: MADKRAHHNA LERKRRDHIK DSFHSLRDSV PSLQGEKASR AQILDKATEY IQYMRRKNHT HQQDIDDLKR QNALLEQQVR ALEGSGC) and Myc^WT^ (MVKRRTHNVL ERQRRNELKR SFFALRDQI PELENNEKAP KVVILKKATA YILSVQAEEQ KLISEEDLLR KRREQLKHKL EQLRNS) were expressed in *E.coli* and purified as previously described (McDuff et al., 2009). The Myc^HEA^ and Myc^RA^ variants, including four additional Myc residues (TEEN) on the the N-terminal side, were expressed in the pET-3a plasmid (Genscript) and purified from the BL21 (DE3) Arabinose-Inducible bacterial strain (Invitrogen) with an adaptation from the same purification protocol (McDuff et al., 2009). Identity and purity of each purified construct was confirmed by mass spectrometry, western blot analysis, SDS-PAGE, and UV spectroscopy.

The double-stranded DNA probe (labeled with the IRD700 fluorophore) and the unlabeled non-specific competitor were ordered from Integrated DNA Technologies (Coralville, IA) and resuspended in DNAse-free water. CACGTG probe: 5’-d(GCG CGG G**<u>CA CGT G</u>**GG CCG GGG)-3’; competitor: 5’-d(GCG CGG G**GG ATC C**GG CCG GGG)-3’. The concentration of the annealed oligonucleotides was confirmed by measuring absorbance at 260 nm. Myc/Max DNA binding was assayed in an Electrophoretic Mobility Shift Assay, as previously described (Beaulieu et al., 2012); briefly, the labeled CACGTG probe was mixed with a 6-fold excess of the unlabeled non-specific competitor, followed by addition of the proteins at a final concentration of 1 μM for each polypeptide (except for the Max only sample: 2 μM) in a final volume of 20 μL. Samples were incubated for 20 minutes prior to loading on the native PAGE in 20 mM Tris-acetate buffer, pH 8.0. Electrophoresis conditions were 100 V for 30 minutes.

### Proliferation assays

For growth curve experiments, 70000 Rat HO15.19 cells were plated in triplicate in 6-well plates and counted every 3 days for 9 days. Similarly, 70000 3T9 cells expressing the various forms of MycER were plated in the presence or absence of 400 nM OHT, and counted every 2 days up to day 6. In the experiments performed with the cb9^Δmyc^ cells, 80000 cells per well were plated in presence of doxycycline for 2 days, then counted and re-plated with or without doxycycline, every two days for the following 10 days. For colony forming assays (CFA), for all cell lines, 10000 cells were plated in 10 cm dishes, let grow for 6-11 days and stained with crystal violet.

For cell cycle analysis, cells were incubated with 33 μM BrdU for 20 min, harvested and washed in PBS, and fixed in ice-cold ethanol. Upon DNA denaturation with 2N HCl for 25 minutes, cells were stained with an anti-BrdU primary antibody (BD Biosciences, #347580) and an FITC-conjugated anti-mouse secondary antibody (Jackson Immunoresearch, # 715-545-150). DNA was stained by resuspending the cells in 2.5 μg/ml Propidium Iodide (Sigma) overnight at 4°C before acquisition with a MACSQuant® Analyzer.

### Antibodies

| **Target** | **Code** | **Company** | **Application** |
| --- | --- | --- | --- |
| c-MYC (Y69) | ab32072 | Abcam | WB |
| Vinculin | V9264 | Sigma | WB |
| FLAG | ab1162 | Abcam | WB |
| HRP-conjugated goat-anti-mouse IgG | 170-6516 | Biorad | WB |
| HRP-conjugated goat-anti-rabbit IgG | 170-6515 | Biorad | WB |
| BrdU | 347580 | Becton Dickinson | FACS |
| c-MYC (N262) | sc-764 | Santa Cruz | ChIP |
| ER-alpha | 06-935 | Merck | ChIP |
| Max | sc-197 | Santa Cruz | WB |

### Genome editing

Bi-allelic deletion of the MYC basic region (BR) as well as insertion of the HEA mutation in the endogenous *c-myc* locus of the cb9 tet-MYC cell line were performed exploiting the type II CRISPR-Cas tool (Ran et al., 2013). Single guide RNA (sgRNA) sequences to target the *MYC* gene in the proximity of the BR-coding region were designed using the online software CRISPR Design Tool (http://crispr.mit.edu/) and cloned as DNA inserts into pSpCas9 (BB)-2A-GFP (PX458) (Addgene plasmid # 48138; a gift from Feng Zhang) (Ran et al., 2013), encoding also the Cas9 protein and GFP. Out of ten tested sgRNAs, we picked the two with the highest cutting efficiency in a surveyor nuclease assay (Ran et al., 2013) (sgRNA7: ACTCCTAGTGATGGAACCC; sgRNA8: ACACGGAGGAAAACGACAAGAGG) Fig. EV3A). We then transfected cb9 cells with sgRNA7 and sgRNA8 together (0.5 µg each), sorted single GFP-positive cells on a 96-well plate, allowed the cells to expand in culture, and screened the resulting cell clones by PCR and Sanger sequencing. We thus obtained one clone, named cb9^Δmyc^, in which both *c-myc* alleles underwent inactivating deletions, although in different ways: one allele encoded a protein missing the BR, Helix I and the loop, and the second a truncated protein lacking the whole C-terminal bHLH-LZ domain (Fig. EV3a).

To generate cb9-*myc*^HEA^ cells, bearing the HEA mutation in both endogenous *c-myc* alleles, the following single stranded donor 192-mer oligonucleotide was used as donor: GCCAAGTTGGACAGTGGCAGGGTCCTGAAGCAGATCAGCAACAACCGCAAGTGCTCGAGTCCCAGGTCCTCAGACACGGAGGAAAACGACAAGAGACGGACAGCCAACGTCTTGGCACGTCAGAGGAGGAACGAGCTGAAGCGCAGCTTTTTTGCCCTGCGTGACCAGATCCCTGAATTGGAAAACAACGAA. We followed the same protocol used above, by co-transfecting the donor, the SpCas9 (BB)-2A-GFP plasmid and sgRNA8, followed by sorting and screening of GFP-positive single-cell clones, yielding a clone with a single *myc*^HEA^ mutant allele. This was subjected to a second round of transfection and clonal selection (with the same donor and plasmid, but a different sgRNA; sgRNA#C1: GACGTTGTGTGTCCGCCTCT), leading to the identification of the homozygous mutant clones cb9-*myc*^HEA^-Cl10 and -Cl33 (Fig. EV3a, b), and a control wild-type clone (cb9-*myc*^WT^-Cl10).

### RNA extraction and RNA-Seq analysis

Total RNA was purified from cell lysates onto Quick-RNA Miniprep columns (Zymo, #R1054) and treated on-column with DNaseI. For RNA-seq experiments, total RNA was purified as above, and RNA quality checked with the Agilent 2100 Bioanalyzer (Agilent Technologies). 0.5-1 μg of RNA were used to prepare libraries for RNA-seq with the TruSeq stranded total RNA Sample Prep Kit (Illumina, #20020596) following the manufacturer’s instructions. RNA-seq libraries were then run on the Agilent 2100 Bioanalyzer (Agilent Technologies) for quantification and quality control and pair-end sequenced on the Illumina 2000 or NovaSeq platforms.

### Single-molecule tracking (SMT) acquisition

The day before single molecule tracking experiments we plated 3T9 cells infected with plasmids expressing HaloTag versions of Myc^WT^, Myc^HEA^ or Myc^RA^ on 4-well LabTek covergrass chambers. One hour before imaging, cells were labeled with 1nM JF549 ligand (Grimm et al., 2016) (Janelia Farm, Ashborn, Virginia, USA), incubated for 30 min at 37°C and extensively washed (two rounds of three washes in PBS followed by 15 min incubation at 37°C in phenol-red free DMEM).

Imaging was carried out on a custom-built microscope capable of inclined illumination (Tokunaga et al., 2008), based on a Olympus IX-73 microscope frame (Olympus Life Science, Segrate, IT), equipped with a stage incubator to control temperature (37°C) and CO_2_ concentration (5%) and a 561nm Diode laser (100mW Cobolt 06-01 series, Cobolt AB, Solna, Sweden), that is synchronized to the camera to achieve stroboscopic illumination. For fast frame-rate acquisitions, we used an Evolve 512 EM-CCD camera (Photometrics, Tucson, AZ, USA), in combination with a ×100, 1.49 NA oil-immersion objective (Olympus Life science), resulting in a pixel size of 158nm. In this case, we set the laser exposure to 2ms, the time between consecutive images *t*_*tl*_ to 10ms and the laser power to ∼ 1 kW cm^−2^ and we collected movies composed by 1000 frames. For slow frame-rate acquisitions, we used a Hamamatsu Orca Fusion sCMOS camera (Hamamatsu Photonics Italia S.r.l, Arese, Italy), combined with a x60, 1.49 NA oil immersion objective (Olympus Life Science), resulting in a pixel size of 108 nm. In this case we set the laser exposure to 50 ms – that results in isolating bound molecules, by motion blurring of the diffusing ones (Chen et al., 2014, Hipp et al., 2019) - the laser power to 100W cm^−2^ and we collected movies composed by up to 200 frames, varying the time between consecutive images *t*_*tl*_ between 200 and 2000 ms. For every experimental condition we acquired at least 30 cells on two experimental days.

### Analysis of the SMT movies – measurement of the bound fraction

The SMT movies collected at fast frame-rate were processed using custom-written Matlab routines, in order to identify and track individual molecules, as previously described (Loffreda et al., 2017, Mazza et al., 2012). A maximum single-molecule displacement of 1.2 □m was allowed between consecutive frames. The resulting tracks were analyzed to quantify the bound fraction, by populating a histogram of single molecule displacements with bin-size Δ*r*, equal to 20nm. The histogram was then normalized in order to provide the probability *p*(*r*)Δ*r* to observe a molecule jumping a distance between *r* − Δ*r*/2 and *r* + Δ*r*/2 in the time between two consecutive frames *t*_*tl*_, which was then fit with a three-component diffusion model, as previously described (Hipp et al., 2019, Loffreda et al., 2017, Speil et al., 2011):

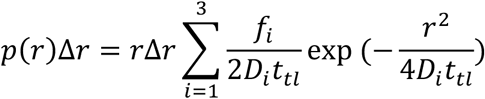

Where *f*_*i*_ is the fraction of molecules moving with a diffusion coefficient equal to *D*_*i*_. Of note for the slowest *D*_1_ we measure diffusion coefficients <0.1μm^2^/s, typical of chromatin bound nuclear proteins at these frame rates (Hansen et al., 2017, Loffreda et al., 2017). *f*_1_ thereby represents the average fraction of bound molecules. To provide standard deviations the fitting parameters a bootstrapping procedure was adopted as described (Hipp et al., 2019). Briefly, we performed multiple fitting iterations, each of them after dropping 20% of the data for each of the data set. Errors are provided as standard deviations of the obtained distribution of parameters following 2000 individual fitting iterations.

### Analysis of the SMT movies – measurement of the residence times

The SMT movies collected at slow frame-rate were processed using the ImageJ plug-in TrackMate (Tinevez et al., 2017). To isolate the bound molecules we allowed a maximum displacement of 220 nm, that allows counting 99% of chromatin-bound molecules (Mazza et al., 2012), and we automatically filled-in gaps of up to three consecutive frames in the tracks. We then computed the cumulative distribution of bound-molecule residence-times and we extracted kinetic parameters on the unbinding process using a global fitting procedure, that allows to minimize the artifacts due to photobleaching, as described (Gebhardt et al., 2013, Hipp et al., 2019). The data was best described by a three-component exponential decay, providing three dissociation constants *k*_1_, *k*_2_, *k*_3_, and the respective weights *F*_1_, *F*_2_, *F*_3_. The average residence time was then calculated as the weighted average: 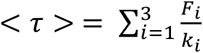. Errors were calculated as SDs from a bootstrapping procedure, as described above.

### Chromatin immunoprecipitation (ChIP)

Cells were washed twice with PBS at room temperature and then fixed for 10 min with formaldehyde 1% in PBS. Fixation was stopped by addition of glycine to a final concentration of 0.125 M. Cells were washed in PBS, scraped in SDS buffer (50 mM Tris at pH 8.1, 0.5% SDS, 100 mM NaCl, 5 mM EDTA, and protease inhibitors) and stored at -80°C before further processing for ChIP as previously described (Sabò et al., 2014). For ChIP-Seq analysis, lysates obtained from 15-30 million cells were immunoprecipitated with 10 μg of antibodies against either Myc (Santa Cruz, #sc-764), ER-alpha (Merck, #06-935). Immunoprecipitated DNA was eluted in TE-2% SDS and crosslinks were reversed by overnight incubation at 65 °C. DNA was then purified on Qiaquick columns (Qiagen) and quantified using QubitTM dsDNA HS Assay kits (Invitrogen). 2-5 ng of ChIP DNA were used for ChIP-seq library preparation as described elsewhere (Blecher-Gonen et al., 2013). ChIP-seq libraries were then run on the Agilent 2100 Bioanalyzer (Agilent Technologies) for quantification and quality control, and sequenced on the Illumina 2000 or NovaSeq platforms. For spike-in controlled Myc ChIP-Seq experiments, we added 5% of HeLa chromatin as a reference exogenous genome to each mouse 3T9 MycER chromatin sample (Bonhoure et al., 2014, Orlando et al., 2014).

### Next generation sequencing data filtering and quality assessment

RNA-seq reads were filtered using the fastq_quality_trimmer and fastq_masker tools of the FASTX-Toolkit suite (http://hannonlab.cshl.edu/fastx_toolkit/). Their quality was evaluated and confirmed using the FastQC application (https://www.bioinformatics.babraham.ac.uk/projects/fastqc/). Pipelines for primary analysis (filtering and alignment to the reference genome of the raw reads) and secondary analysis (expression quantification, differential gene expression) have been integrated in the HTS-flow system (Bianchi et al., 2016). Bioinformatic and statistical analyses were performed using R with Bioconductor and comEpiTools packages (Gentleman et al., 2004, Kishore et al., 2015).

### ChIP-seq data analysis

ChIP-Seq NGS reads were aligned to the mouse reference genome mm9 (or human hg19 for spike-in controlled Myc ChIP-Seq), through the BWA aligner^11^ using default settings. Before comparison of ChIP-seq samples, ambiguous reads mapping to both mm9 and hg19 were identified with Picard tools and removed. Peaks were called using the MACS2 software (v2.0.10)^12^ with the option ‘– mfold = 7,30 -p 0.00001 -f BAMPE’, thus outputting only enriched regions with P-value <10-5. Promoter peaks were defined as all peaks with at least one base pair overlapping with the interval between - 2 kb to +2 kb from the nearest TSS. The presence of canonical and variant E-boxes (CACGCG, CATGCG, CACGAG, CATGTG)^13-15^ in Myc ChIP-seq peaks was scored in a region of 100 bp around the peak summit. When comparing Myc ChIP-seq with H3K4me3, H3K4me1 or H3K27ac histone marks to define peaks in active promoter or enhancers^16,17^, we considered two peaks as overlapping when sharing at least one base pair (findOverlaps tool of the comEpiTools R package). Motif discovery was performed using MEME-ChIP suite (Bailey et al., 2009) with default parameters using as input the regions ± 100 bp around peak summits reported by MACS2. For heatmap and intensity plots, we used bamCoverage from deepTools 3.3.1 (Ramirez et al., 2016) to calculate read coverage per 10-bp bin using RPKM normalization option. For spike-in controlled Myc ChIP-Seq samples, the normalization scaling factor was calculated as previously described (Orlando et al., 2014), using the option –scaleFactor. Heatmaps were performed through the functions computeMatrix followed by plotHeatmap from deepTools using the normalized bigwig files.

### RNA-seq data analysis

RNA-seq NGS reads were aligned to the mm9 mouse reference genome using the TopHat aligner (version 2.0.8) (Kim et al., 2013) with default parameters. In case of duplicated reads, only one read was kept. Read counts were associated to each gene (based on UCSC-derived mm9 GTF gene annotations), using the featureCounts software (http://bioinf.wehi.edu.au/featureCounts/) (Liao et al., 2014) setting the options -T 2 -p -P. Absolute gene expression was defined determining reads per kilobase per million mapped reads defining total library size as the number of reads mapping to exons only (eRPKM). Differentially expressed genes (DEGs) were identified using the Bioconductor Deseq2 package (Love et al., 2014) as genes whose q-value is lower than 0.05. Functional annotation analysis to determine enriched Gene Ontology categories was performed using the online tool at http://software.broadinstitute.org/gsea/index.jsp. Gene set enrichment analysis (GSEA) was performed using the Desktop tool of the Broad Institute (http://software.broadinstitute.org/gsea/index.jsp) with custom gene lists (Tesi et al., 2019).

### Statistical analysis

All experiments were performed at least in biological triplicates. Sample size was not predetermined, but is reported in the respective Figure legends. Two-tailed Student’s t-test was used to compare between two groups and expressed as p-values.

## Acknowledgements

We thank Stefano Campaner, Gioacchino Natoli, Sebastiano Pasqualato, Matteo J. Marzi, Andrew Wilkie and colleagues in our group for discussion, insight and suggestions. We also thank L. Rotta, S. Bianchi, T. Capra and L. Massimiliano for assistance with Illumina sequencing. LS, VCC and MEB acknowledge the Cellex Foundation for providing research facilities and equipment. AL and DM were supported by Fondazione Cariplo (GR: 2014-1157) and by the Italian Association for Cancer Research (AIRC, IG 2018-21897). Work in the Amati lab was supported by funds from the European Research Council (grant agreement no. 268671-MycNEXT), the Italian Health Ministry (RF-2011-02346976) and AIRC (IG 2015-16768 and IG 2018-21594).

## Author contributions

PP, TK and AS designed and performed most of the experiments described in this work. AV and MD provided technical support. MD, MF, MJM and AS performed bioinformatic data analysis. AL and DM produced and analysed data with single-molecule tracking microscopy. MEB, VCC and LS produced the *in vitro* binding data. MM contributed to the structure-based design of the Myc mutants. BA and AS conceived the project, co-supervised the work, and wrote the manuscript.

## Data availability

The RNA-Seq and ChIP-Seq data described in this work are accessible through Gene Expression Omnibus (GEO) series accession number GSE147639; previously described ChIP-seq data (H3K4me3, H3K4me1, H3K27ac, Dnase I, RNAPII) (Sabò et al., 2014) are accessible through the accession number GSE51011. All R scripts used in data analysis and generation of Figures are available upon request.

## Figure Legends

**Expanded View Figure 1.**
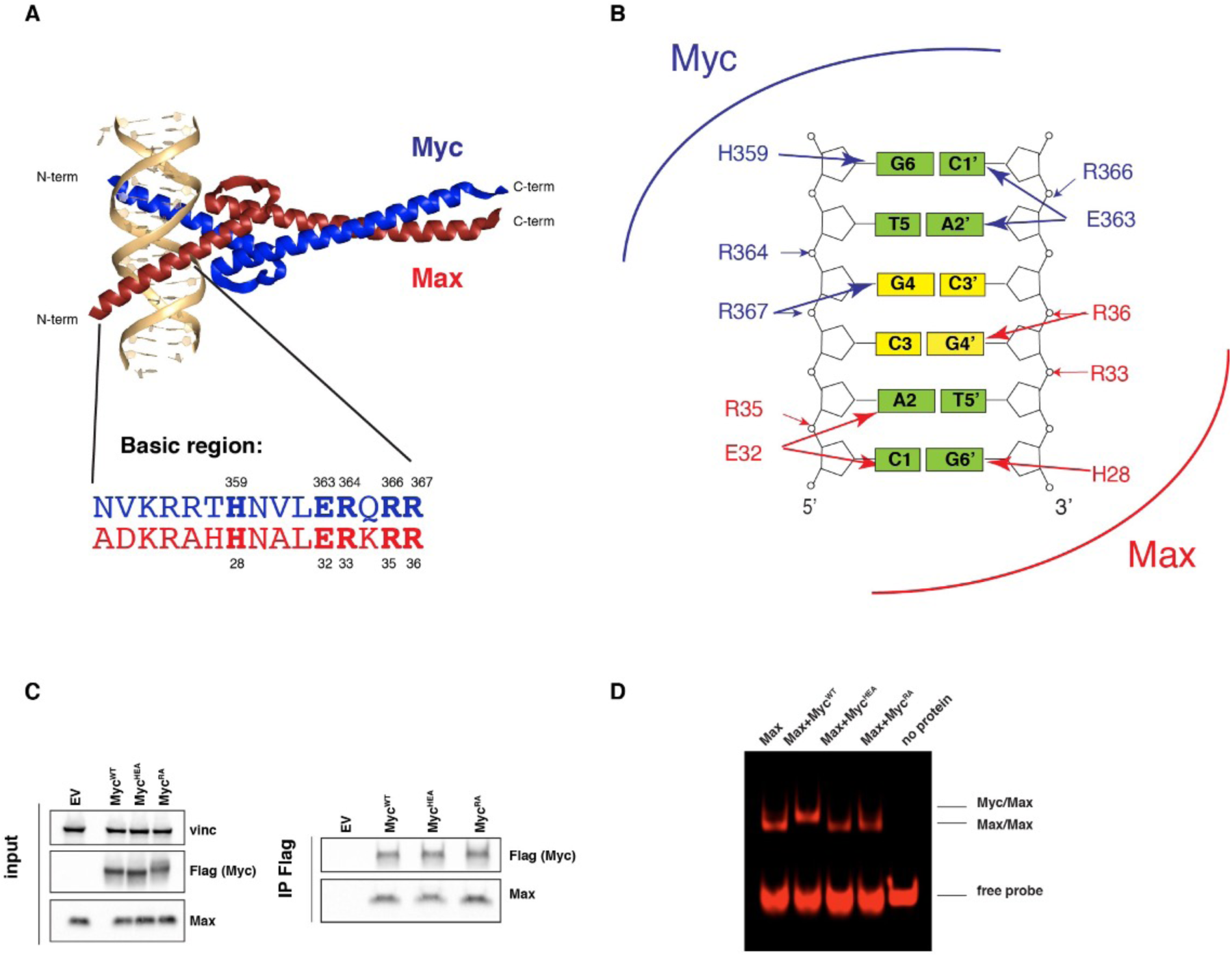
Design and biochemical characterization of Myc mutants. **A** Structure of the DNA-bound Myc/Max dimer (Nair & Burley, 2003), with an alignment of the Myc and Max basic regions (numbering based on the 439 a.a. human MYC protein (GenBank nr. AAA36340.1). **B** H359 and E363 establish H-bonds with the complementary G6 and C1’, bases, respectively, E363 forming an additional bond with A2. Note that, in keeping with the symmetric configuration of the Myc/Max dimer (panel a) and with the similar structure of the Max homodimer (Ferre-D’Amare et al., 1993), the corresponding residues in Max (R33/35/36, and H28/E32) form equivalent contacts with the other half of the E-box palindrome. **C** Immunoblot analysis of 293T cells transfected with plasmids expressing the indicated proteins (left). Lysates were immunoprecipitated (IP) in with anti-Flag beads, and the precipitates subsequently analyzed by immunoblotting (right) with the indicated antibodies. **D** Electrophoretic Mobility Shift Assay with the bHLH-LZ of Max, alone or in a 1:1 molar ratio with either of the Myc bHLH-LZ peptides (Myc^WT^, Myc^HEA^ or Myc^RA^). The polypeptides purified from *E.coli* were incubated at a 1:1 molar ratio with a fluorescently-labelled CACGTG DNA probe, and the complexes separated on a native polyacrylamide gel, as described (Beaulieu et al., 2012).

**Expanded View Figure 2.**
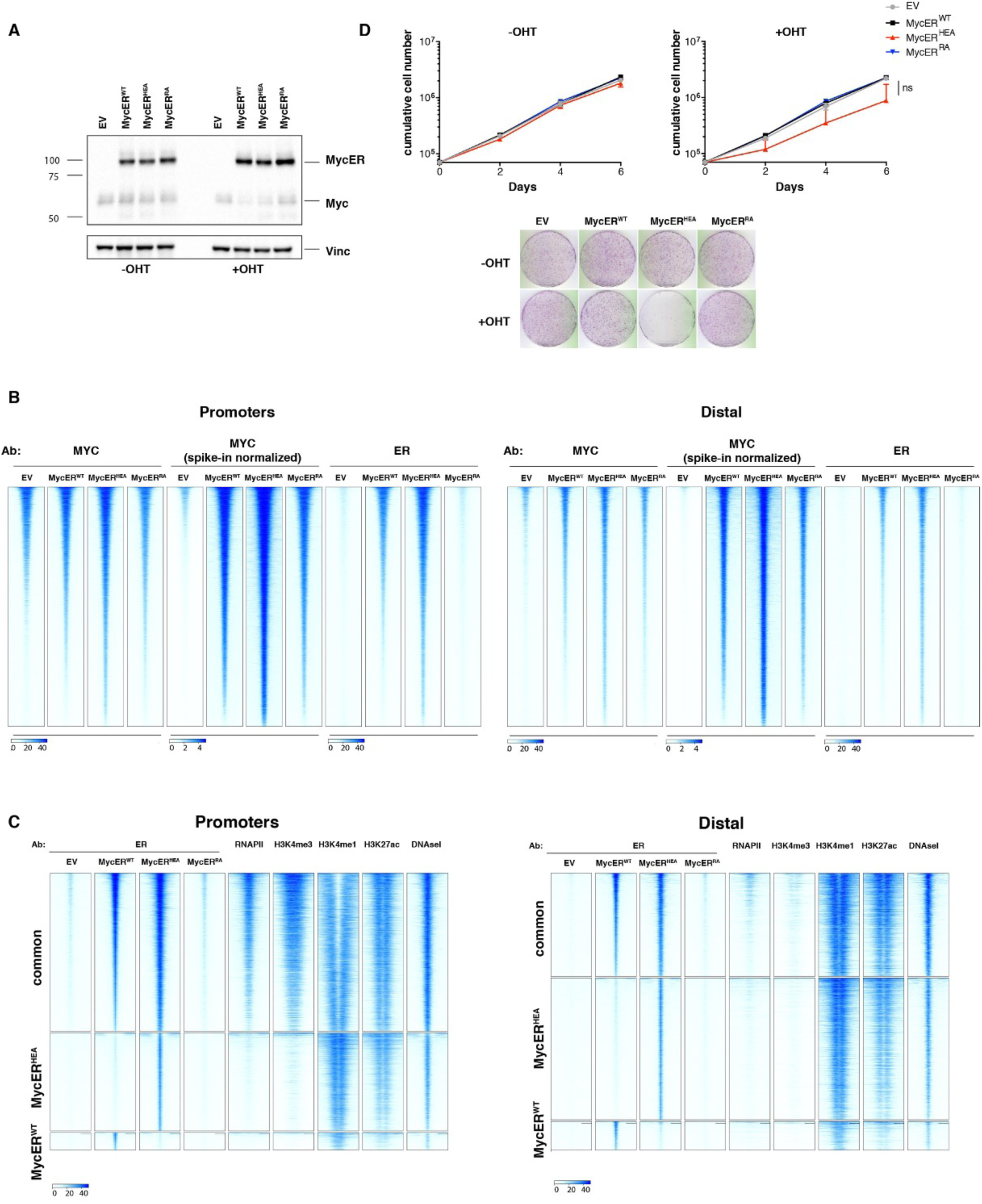
Characterization of 3T9 MycER^WT^, MycER^HEA^, MycER^RA^ - infected cells. **A** Immunoblot analysis of 3T9 cells infected with retroviral vectors expressing the indicated MycER proteins, treated or not with OHT (4h), as indicated. **B** Heatmaps representing normalized ChIP-seq intensities at Myc or MycER-associated promoters or distal sites, as indicated. Each row represents a genomic site called in at least one of the experimental samples, with each column spanning a 4 kb-wide genomic interval centered on the union of MycER peaks (called either with anti-Myc or anti-ER antibody). The Myc ChIP-Seq samples are shown before and after spike-in normalization with human chromatin. All rows were ranked on the basis of the signal intensity (without spike-in normalization) of the MycER^WT^ sample. **C** Heatmaps representing normalized ChIP-seq intensities at MycER^WT^- and/or MycER^HEA^-associated promoters or distal sites. Each row represents a 4 kb-wide genomic interval centered on the Myc peak summit in either the MycER^WT^ sample (for the common and unique MycER^WT^ sites) or the MycER^HEA^ sample (for its unique sites). All rows were ranked on the basis of the signal intensity of the MycER^WT^ sample (considering all signals, regardless of peak calling). **D** Growth curves and colony forming assays for 3T9 MycER expressing cells in presence or absence of OHT.

**Expanded View Figure 3.**
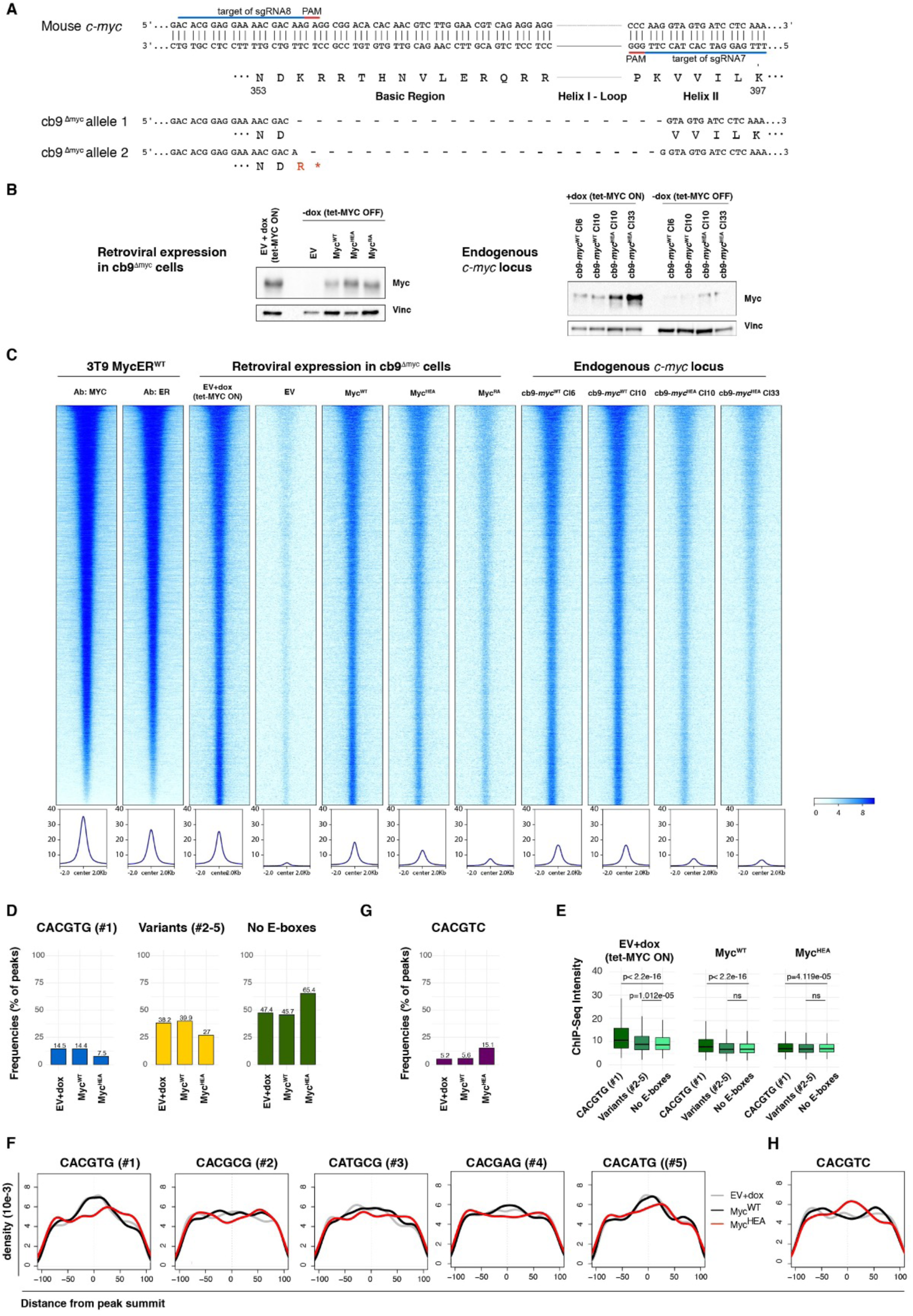
Myc^HEA^ retains widespread non-specific DNA binding, but loses E-box recognition. **A** Schematic representation of the mutant *c-myc* alleles in cb9^Δmyc^ cells (numbering based on the 439 a.a. mouse Myc protein: NCBI nr. NP 001170823.1). **B** Immunoblot analysis of cb9^Δmyc^ cells infected with retroviral vectors expressing the indicated Myc proteins (left) and cb9-*myc*^HEA^ cells and control cb9-*myc*^WT^ clones (right), cultured with or without doxycycline, as indicated. **C** Heatmaps representing normalized Myc ChIP-seq intensities in cb9^Δmyc^, cb9-*myc*^HEA^ and control cells, as indicated. As a reference, the MycER^WT^ profiles obtained with anti-Myc or anti-ER antibodies (Ab) are also shown. Each row represents a genomic site called in at least one of the experimental samples, with each column spanning a 4 kb-wide genomic interval centered on the union of MycER peaks. All sites are ranked according to the intensity of the MycER^WT^ ChIP signal performed with the anti ER antibody. The graphs below the heatmaps shows normalized intensity profiles for each sample. **D** As Fig. 1B: Frequency of peaks (as %) that contain the indicated motif (within ±100 bp from the peak summit) in each ChIP-seq sample (EV, WT, HEA and RA). **E** As Fig. 1C: Average ChIP-seq intensities for regions containing the indicated motifs in each sample. (**F**) As Fig. 1D: density plots showing the distribution of the indicated motifs in a region of ±100 bp from peak summit. **G, H** as in (D) and (F), respectively, for the CACGTC motif.

**Expanded View Figure 4.**
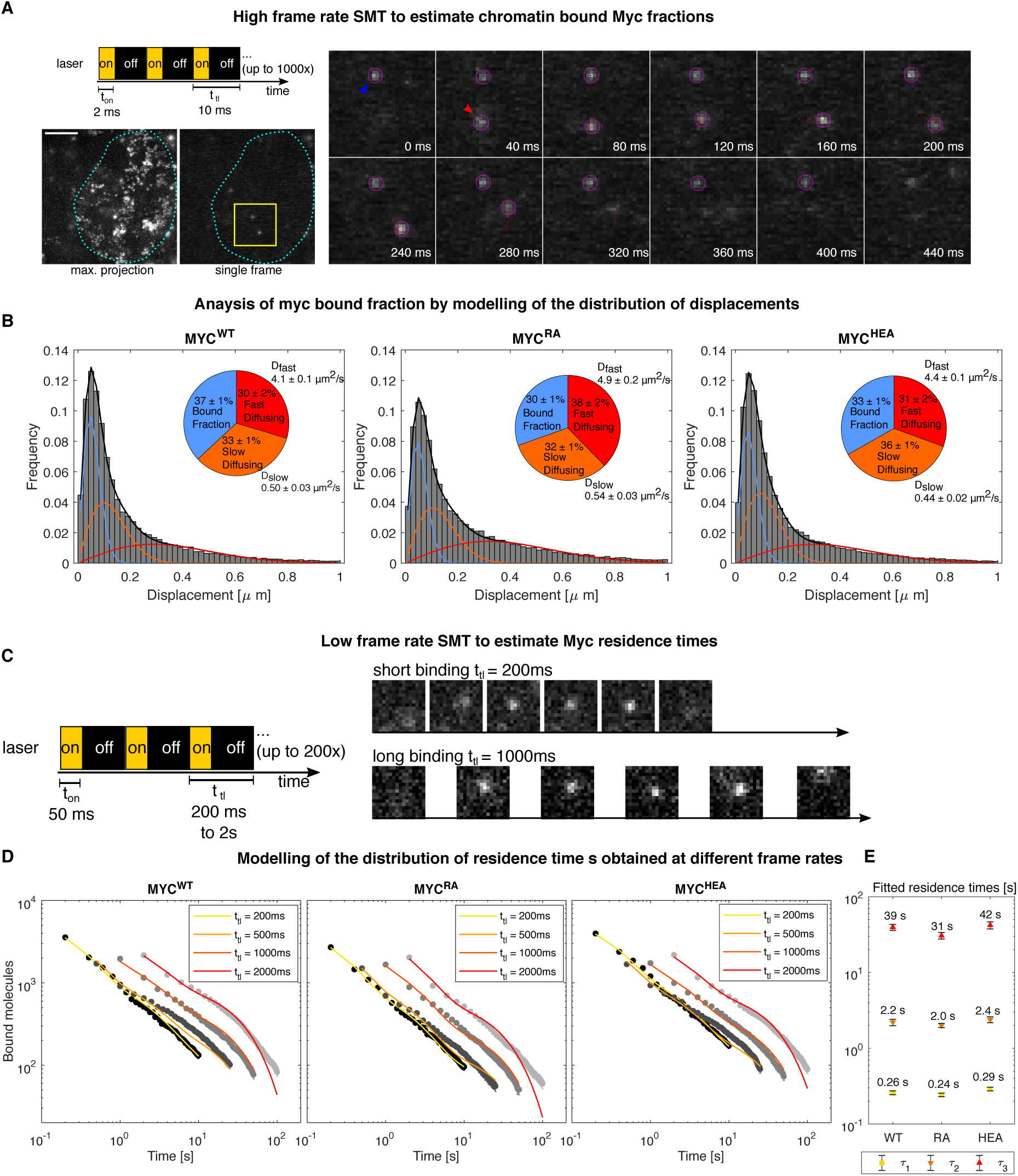
Single molecule microscopy analysis of Myc mutants. **A** Schematic representation of the illumination protocol for the SMT acquisitions to quantify the fraction of bound Myc molecules, and exemplary acquisitions. The maximal projection of a representative Myc^WT^ movie is shown, displaying the nucleus boundaries (cyan dotted line) and a representative region (yellow square), for which exemplary frames are displayed on the right. The blue and red arrows highlight a bound and a diffusing molecule, respectively. Scale-bar: 5 µm. **B** Tracking the single Myc molecules allows estimating the distribution of displacements which is fit with a three-component diffusion model to extract the fraction of bound molecules as well as the fractions and the diffusion coefficients of the diffusing molecules (see Methods). **C** Schematic of the illumination protocol for the SMT movies to quantify the residence times of bound Myc molecules. Acquisition at different frame rates (*t*_*tl*_ ranging between 200 ms and 2s) are performed to measure the residence times of bound Myc molecules at multiple time-scales and to correct for photobleaching. **D** The cumulative distributions of residence times are analyzed together using a global model accounting for photobleaching. **E** The model allows to estimate three characteristic times *τ*_1_, *τ*_2_, *τ*_3_ – inverse of the three characteristic reaction rates, than are then averaged together to provide an estimate of the average residence time of the various Myc proteins on chromatin (See Methods and Fig. 1F).

**Expanded View Figure 5.**
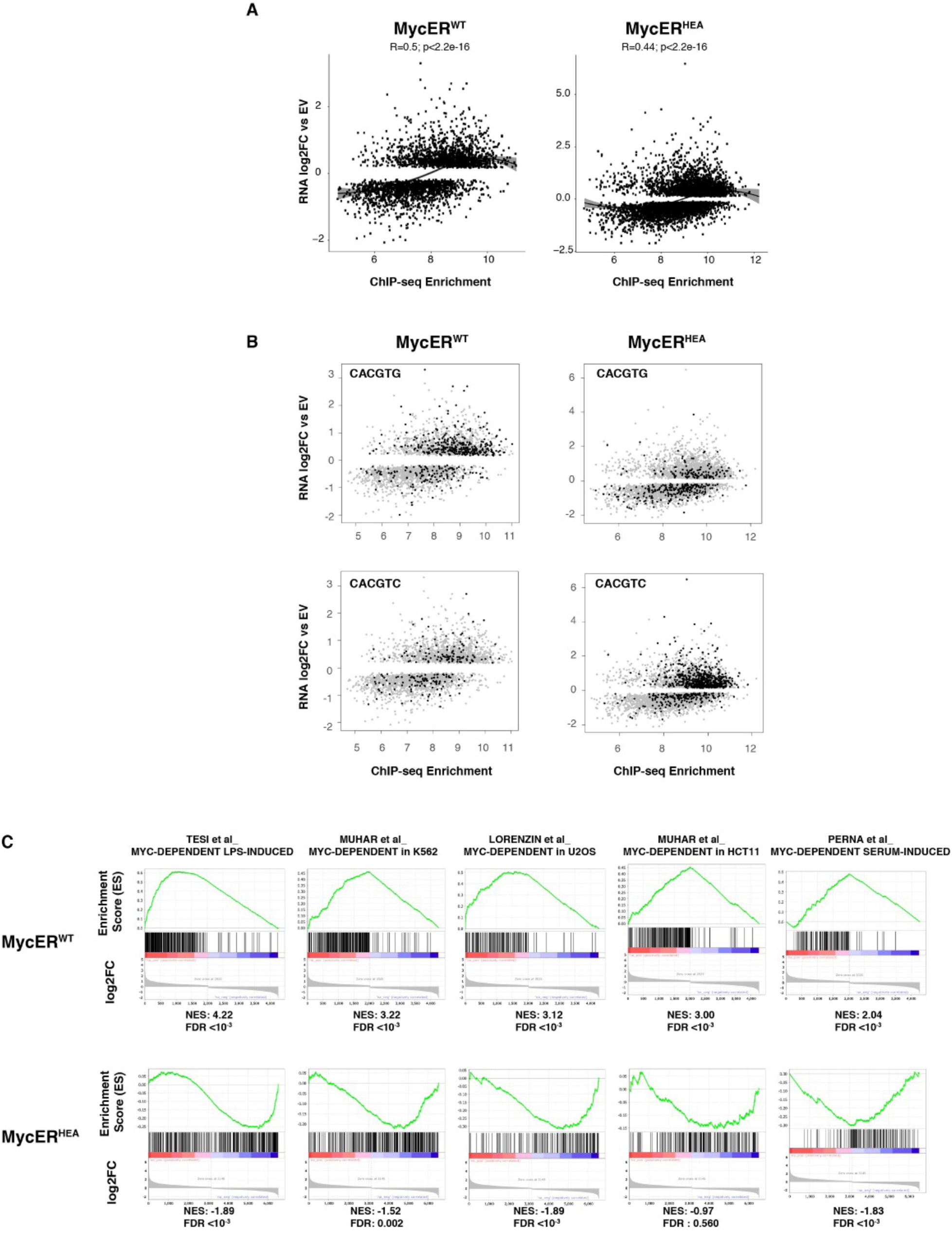
Transcriptional analysis of MycER proteins. **A** Scatter plot showing ChIP-seq enrichment with anti-Myc antibodies (x-axis) and the RNA-seq log2 fold change (log2FC) relative to EV (y-axis) for all DEGs (qval<0.05) with a peak on the promoter in MycER^WT^ (left) or MycER^HEA^ (right). **B** as in (A), with black dots indicating the presence of the indicated motif within ±100bp from the peak summit. **C** Gene set enrichment plots for 5 custom Myc-dependent signatures (see Methods for details and references). Normalized Enrichment score (NES) and False Discovery Rate (FDR) values) are reported for each dataset. Genes were sorted from left to right according to the log2FC in their expression when comparing MycER^WT^ or MycER^HEA^ versus empty vector.

**Appendix Figure S1.**
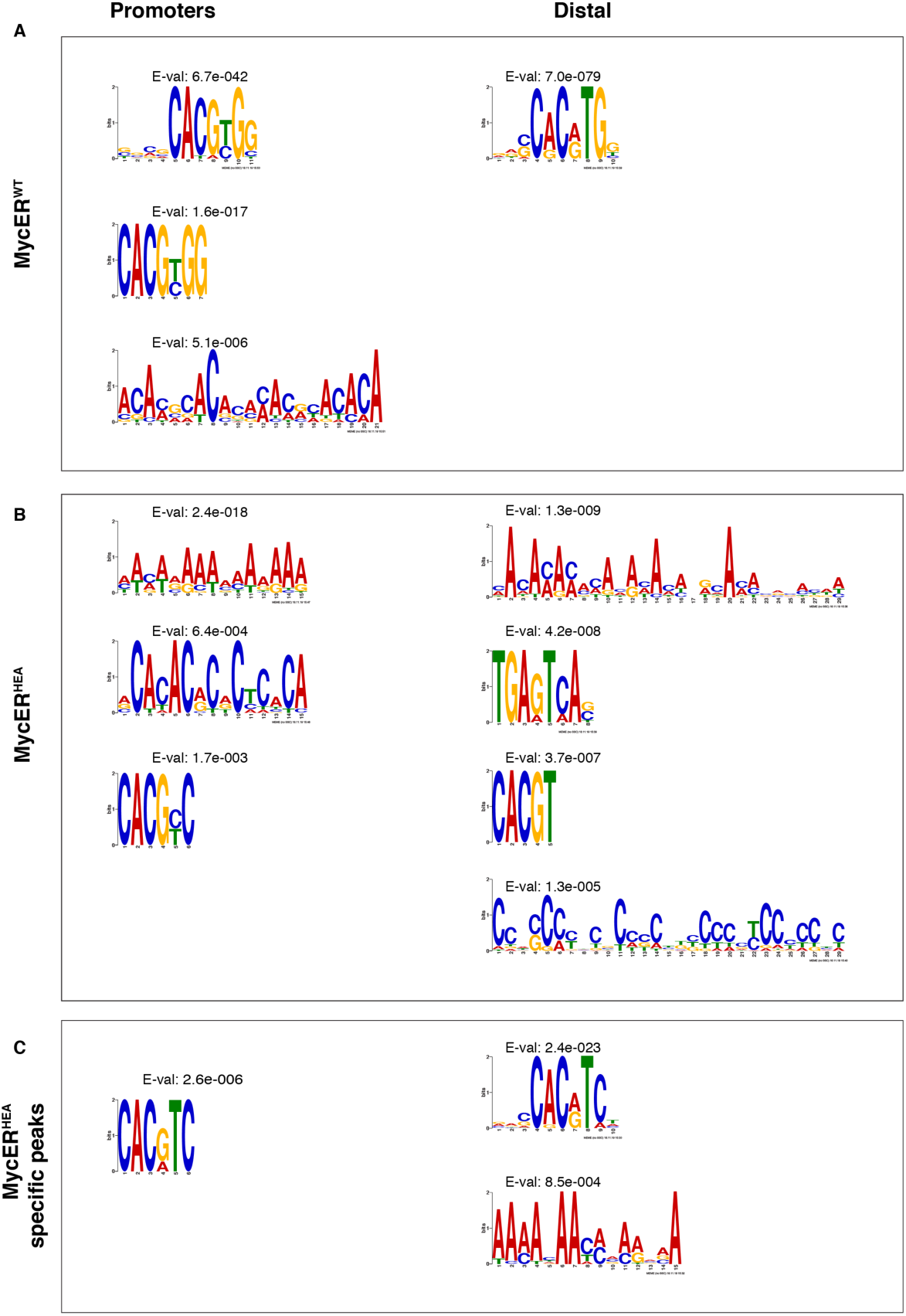
*De novo* motif discovery analysis of Myc mutants binding sites at promoter and distal regions. **A** *De novo* motif discovery analysis performed underneath the summit of the top 200 MycER^WT^ promoter-associated or distal MycER^WT^ peaks. Position weight matrixes of predicted DNA binding motifs are shown together with their E-values. **B** As in (A) for the top 200 MycER^HEA^ peaks. **C** As in (A) for MycER^HEA^ -specific peaks (i.e. not bound by MycER^WT^).

